# RUNX1 is Expressed in a Subpopulation of Dermal Fibroblasts and Higher RUNX1 Levels are Associated with the Severity of Systemic Sclerosis

**DOI:** 10.1101/2024.04.03.587966

**Authors:** Rezvan Parvizi, Zhiyun Gong, Helen C. Jarnagin, Diana M. Toledo, Tamar R. Abel, Dillon Popovich, Madeline J. Morrisson, Sasha Shenk, Monique E. Hinchcliff, Jonathan A. Garlick, Patricia A. Pioli, Michael L. Whitfield

## Abstract

The activation of Runt-related transcription factor 1 (RUNX1) in fibroblasts has been implicated in wound healing and fibrosis; however, the role of RUNX1 in the fibrotic progression of the autoimmune disease systemic sclerosis (SSc) is not known. Through gene expression analysis, we have demonstrated an association between the severity of dermal fibrosis and the expression levels of *RUNX1* in the skin of patients with SSc. Additionally, we identified hypomethylated CpG sites proximal to the *RUNX1* gene, implicating their potential role in the increased expression of *RUNX1*. Analysis of single-cell RNA-seq data from skin biopsies of individuals with SSc revealed that *RUNX1* is higher in subpopulations of fibroblasts enriched in SSc, which are believed to contribute to fibrosis. Lastly, modulation of *RUNX1* activity using an inhibitor caused a reduction in fibroblast contraction and proliferation rates. Altogether, this study is the first to demonstrate a potential role for *RUNX1* in the pathogenesis of systemic sclerosis dermal fibrosis.

## Introduction

Dermal fibrosis is a major clinical manifestation of systemic sclerosis (SSc). It results from the uncontrolled deposition of extracellular matrix (ECM) by fibroblasts residing in the skin^1^. A physician-performed assessment of the modified Rodnan Skin Score (mRSS) is commonly used to evaluate the degree and progression of dermal fibrosis, and severity of SSc disease. There are two clinical types of the disease, based on the extent of skin involvement: limited cutaneous SSc (lcSSc) and diffuse cutaneous SSc (dcSSc)^2^. We have developed a molecular classification system, termed the ‘intrinsic’ subtypes, which is based on gene expression changes observed in lesional and non-lesional skin biopsies. This classification categorizes patients with SSc into four subtypes: inflammatory, fibroproliferative, normal-like, and limited^3^. As dermal fibrosis progresses, patients with SSc experience skin thickening, distortion, and tightening that can limit joint movement and cause significant discomfort^4^. The collective cross-talk between dermal transcriptional patterns and signaling pathways contributes to fibrotic activation in SSc^5^. Through network analyses of transcription factor (TF) activity, we have previously shown that the *runt-related transcription factor 1* (*RUNX1*) is a key regulator in SSc. *RUNX1* is highly expressed in SSc skin biopsies, particularly within the inflammatory patient subgroup, and is strongly correlated with measures of disease severity^6^.

RUNX1 is a member of a family of DNA-binding transcription factors that partners with a constitutively expressed core-binding factor subunit-β (CBFβ) to form an active heterodimer that regulates the expression of downstream genes. The *RUNX1* gene is located on chromosome 21 and encodes for three isoforms that differ in their N-terminal coding sequences^7^. Transcriptional, translational, and post-translational modifications tightly regulate the expression of *RUNX1*. Significantly, several studies have documented crosstalk between RUNX1 and profibrotic signaling pathways, including transforming growth factor-β (TGF-β)^8^, NF-κB^9^, and Wnt signaling^10^. While the importance of RUNX1 in most hematopoietic cell lineages has been characterized in detail, its role in dermal fibroblasts and fibrosis is not clear.

*RUNX1* is overexpressed in a range of human cancers^11–13^, as well as autoimmune^14^ and fibrotic diseases^15,16^. It was also recently identified as being actively involved in wound healing processes^17^. A spatial multiomic analysis of wound healing has demonstrated that RUNX1 is highly active in inner wound mechano-fibrotic fibroblasts, which differentiate into a strongly profibrotic cellular subset^17^. Moreover, expression of *RUNX1* is significantly increased in human glioma tissues and is closely associated with tumor grade and the expression of ECM-related genes such as *Fibronectin 1* (*FN1*), *collagen type IV alpha 1 chain* (*COL4A1*), and *Lumican* (*LUM*), which are all implicated in the pathogenesis of glioblastoma^18^. *RUNX1* promotes the development of pulmonary arterial hypertension^19^. It is also more highly expressed in the border zone of cardiac remodeling that occurs after myocardial infarction; when targeted in that case, RUNX1 inhibition has a protective effect and preserves cardiac contractile function^16,20,21^. Study of synovial biopsies from patients with rheumatoid arthritis and plaque biopsies from patients with atherosclerosis also identified *RUNX1* as a top overlapping key transcription factor^14^. Although *RUNX1* has been implicated as a contributor to pathology in fibrotic disease, the role of *RUNX1* in the pathogenesis of SSc dermal fibrosis has not yet been determined.

In this study, we present the first evidence demonstrating the involvement of *RUNX1* expression in a subpopulation of dermal SSc fibroblasts, and also investigate its significance to SSc disease severity. Using publicly available skin transcriptional datasets and analyzing data from skin of 200 patients with SSc, we characterized *RUNX1* expression patterns and investigated its epigenetic profile to establish a comprehensive understanding of its role in SSc dermal fibroblasts.

## Results

### *RUNX1* expression is significantly higher in the skin of diffused SSc patients and is associated with a TGF-β fibroblast gene signature

Our prior work indicated a role for the *RUNX1* transcription factor in the inflammatory subtype of SSc^6^. We began this study by analyzing a larger cohort of skin biopsies from individuals with SSc and healthy controls to examine the distribution of *RUNX1* expression and to further investigate its correlation with disease severity^6^. Analysis of gene expression data showed significantly increased expression of *RUNX1* in lesional forearm skin biopsies from 86 individuals diagnosed with lcSSc and dcSSc^22^. *RUNX1* expression was highest among individuals diagnosed with early dcSSc and decreased over a course of three years (**Fig. 1A**, 1B). *RUNX1* was also elevated in samples of non-lesional skin, i.e., samples collected from the flank, of patients with SSc, compared to healthy control non-lesional skin (Supp **Fig. 1**), suggesting that the disease-specific activation of *RUNX1* is shared between early fibrotic skin and pre-lesional/pre-fibrotic skin biopsies. Increased expression of *RUNX1* was subsequently confirmed in five additional publicly available datasets (GSE9285, GSE32413, GSE125362, GSE97248, and GSE58095), totaling more than 120 individuals with SSc ^3,23–27^ (Supp **Fig. 2A**-**2E**, Supp Table S1). At baseline, individuals with early disease (defined as biopsies taken from less than two years since first onset of non-Raynaud’s symptoms) exhibited higher levels of *RUNX1* expression compared with those with a disease duration of greater than two years (**Fig. 1C**). A significant positive correlation was found between *RUNX1* expression and mRSS in two separate cohorts (*r^2^* = 0.31, *P* = 1.3e-06; *r^2^* = 0.25, *P* = 4.9e-05)^25,27^ (**Fig. 1D**, Supp **Fig. 2F**). *RUNX1* was highest among SSc patients with a higher skin score at the site of biopsy (Supp **Fig. 2G**) and those with ILD (Supp **Fig. 2H**), consistent with a prior report^6^, indicating a potential association between disease severity and *RUNX1* expression levels.

**Figure 1.**
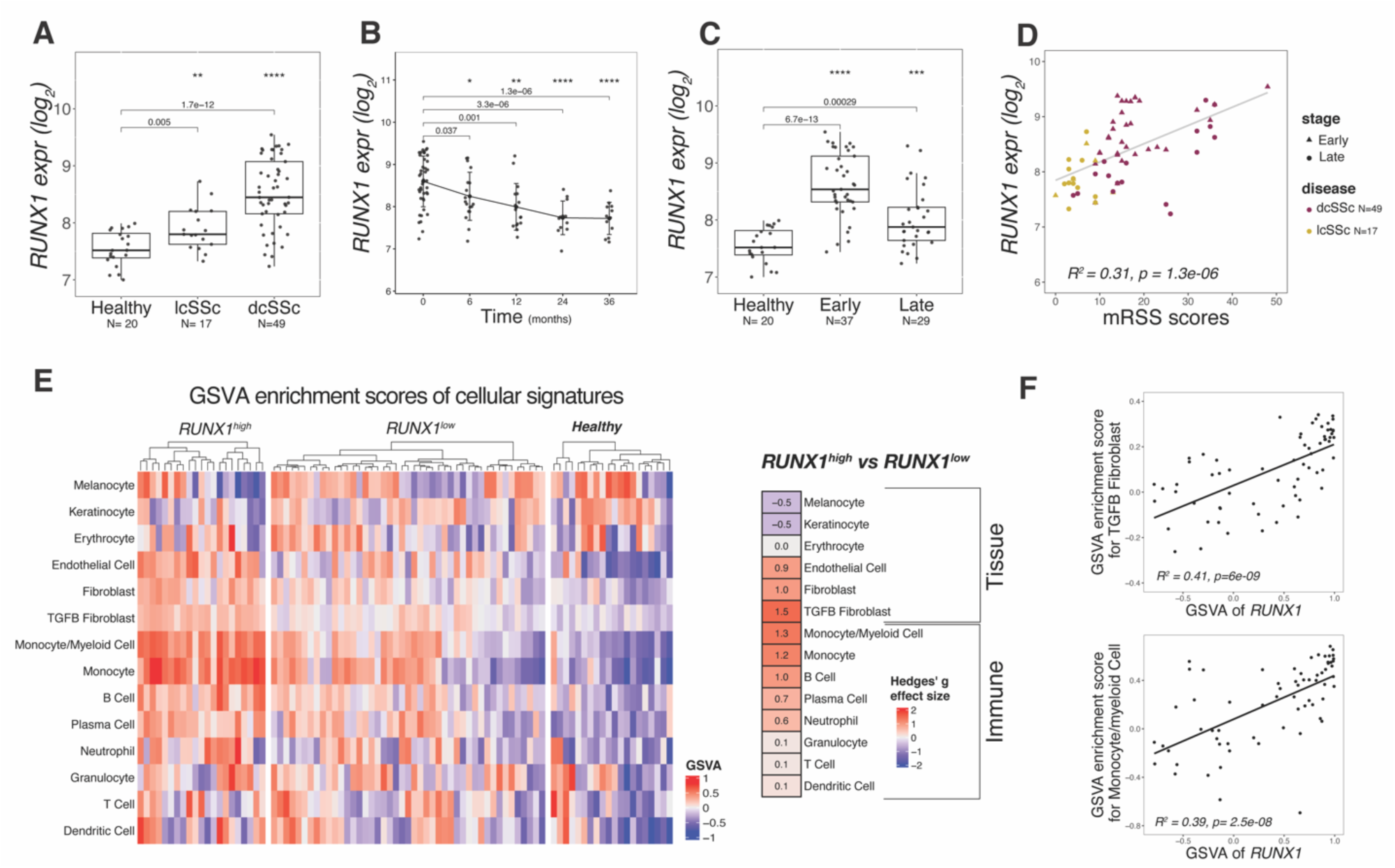
*RUNX1* expression is significantly higher in the skin of patients with dcSSc and is associated with TGF-β fibroblast and myeloid cell signatures. **(A)** *RUNX1* expression rate in forearm skin biopsies of dcSSc (*N*=49), lcSSc (*N*=17), and healthy (*N*=20) patients. **(B)** The expression rate of *RUNX1* over the course of three years (data presented for 0, 6, 12, 24, and 36 months). (C) *RUNX1* expression rate for healthy and SSc patients at early or late stages of disease at baseline. **(D)** Correlation between *RUNX1* expression and mRSS skin score at baseline for both lcSSc (yellow) and dcSSc (red) (*N*=66); early-stage patients are shown as a triangle and late-stage patients as a circle. **(E)** GSVA enrichment scores of main cellular signatures in healthy (*N*=20), patients with *RUNX1*^high^ (*N*=21) and *RUNX1*^low^ (*N*=45). Hedge’s g effect size of *RUNX1*^high^ vs. *RUNX1*^low^ is presented in the graph. **(F)** Correlation between the TGF-β fibroblast and monocyte and myeloid cell signatures with *RUNX1*.

**Figure 2.**
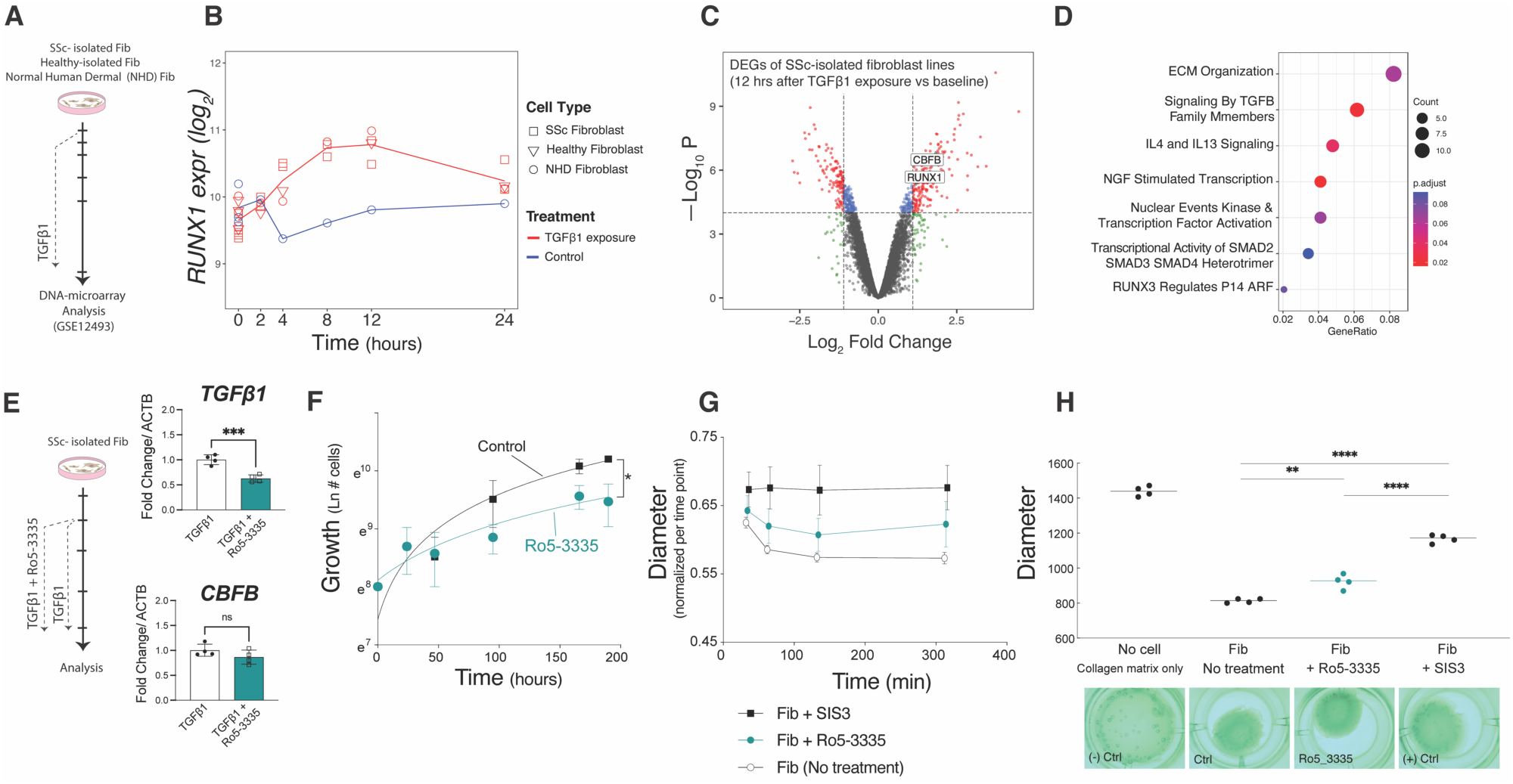
TGF-β1 increases the *RUNX1* expression in SSc fibroblasts and inhibition of *RUNX1* constrains fibroblast proliferation and contraction. **(A)** Schematic graph illustrating the timeline for the culture and TGF-β1 treatment of dcSSc-isolated fibroblasts, matched healthy-isolated fibroblasts, and Normal Human Dermal (NHD) fibroblast cells. (B) *RUNX1* expression rate in samples treated with TGF-β1 (in red) vs. control for the 24 hours after exposure. **(C)** Volcano plot of differentially expressed analysis of the two SSc-isolated fibroblast lines at 12 hours after exposure vs. the baseline. **(D)** The pathway analysis of Reactome gene sets shows the biological pathways and processes that are significantly represented within top DEG genes of SSc-isolated fibroblast lines 12 hours after TGF-β1 treatment vs. the baseline. **(E)** Fold change expression of *TGF-β1* and *CBFB* in TGF-β1-induced SSc fibroblasts treated with Ro5-3335 (*RUNX1* inhibitor), compared to control. **(F)** Proliferation curve of NHD fibroblasts in the presence and absence of Ro5-3335. **(G–H)** The 3D collagen contraction assays, fixed **(G)** and floating (**H**) models, of NHD fibroblasts treated with Ro5-3335. SIS3 (SMAD3 inhibitor) was used as positive control that significantly eliminates the contraction ability of fibroblasts. (Student’s t-test *P-*value: **0.001–0.01, ****<0.0001 in GraphPad Prism v9)

We next set out to identify the cell-type-specific signaling pathways with which *RUNX1* expression was most closely associated. We performed gene set variation analysis (GSVA) to calculate the enrichment scores^28^ of fibroblasts, endothelial cells, keratinocytes, and immune cells for 86 individuals at baseline (**Supp Table S2**). SSc patients with a high mRSS score (more than 10) were divided into two groups: *RUNX1*^high^, defined as *RUNX1* expression over 1 standard deviation (SD) above the mean, and *RUNX1*^low^. The GSVA scores with the strongest enrichment in *RUNX1*^high^ SSc patients were the TGF-β-activated fibroblast and monocyte/myeloid cell gene signatures (**Fig. 1E**). Hedge’s g effect size was used to indicate the magnitude of the difference between groups and confirmed strong enrichment of these pathways (effect sizes of 1.5 and 1.3, respectively). Similarly, evaluation of the Pearson’s correlation between the major cellular signatures and *RUNX1* demonstrated that TGF-β-activated fibroblast (*r^2^* = 0.41, *P* = 6e-09) and monocytes/myeloid cell gene signatures (*r^2^* = 0.39, *P* = 2.5e-08) had the highest correlations with *RUNX1* mRNA expression (**Fig. 1F**).

### RUNX1 regulation is associated with the activation of TGF-β1 signaling in SSc fibroblasts

To assess the regulation of *RUNX1* by the TGF-β1 signaling pathway, we investigated the expression of *RUNX1* in fibroblasts after exposure to exogenous TGF-β1. We analyzed a DNA microarray dataset previously generated by our lab, consisting of two independent SSc fibroblasts, one healthy control fibroblast isolated in parallel, and one normal human dermal (NHD) fibroblast cell lines obtained from ATCC. All treated with 50 pM of TGF-β1^29^ (**Fig. 2A**, available at the NCBI GEO GSE12493). As shown in **Fig. 2B**, *RUNX1* expression was significantly induced by TGF-β1 (relative to vehicle alone treatment) as soon as 4 hours after treatment and remained elevated until our experimental endpoint, at 24 hours. Differential gene expression analysis, comparing SSc fibroblasts collected after 12 hours of TGF-β1 treatment with baseline, showed significantly elevated levels of *RUNX1* and *CBFB* mRNA (**Fig. 2C**). Analysis of the differentially-expressed pathways using Reactome showed that the top pathways that were associated with high *RUNX1* expression in SSc fibroblasts after TGF-β1 exposure related to “ECM organization”, “signaling by TGF-β family members”, and “IL4 and IL13 signaling” (**Fig. 2D**).

To investigate whether the inhibition of TGF-β would reduce *RUNX1* expression in SSc skin, we analyzed data from a clinical trial of fresolimumab, conducted by Rice *et al.* in 2015. Fresolimumab is a human monoclonal antibody that binds to and inhibits all three isoforms of TGF-β (TGF-β1–3). In this study, participants were divided into two treatment arms: one that received two low-dose infusions of fresolimumab and another that received one high-dose infusion of fresolimumab. Mid-forearm skin biopsies were collected at baseline and at 3, 7, and 24 weeks after treatment. While no significant changes were observed in samples analyzed at 24 weeks post-treatment, expression of *Thrombospondin 1 (THBS1)*, which is used as a biomarker of TGF-β pathway activation, was decreased 3 and 7 weeks after treatment in patients that received high-dose fresolimumab. We re-analyzed these data and specifically interrogated the expression of *RUNX1,* which was decreased by fresolimumab treatment, paralleling findings with *THBS1* (**Supp Fig. 3A**). Analysis of the TGF-β fibroblast gene signature at baseline and three weeks after a single high-dose treatment with fresolimumab showed that TGF-β-associated genes had decreased expression (**Supp Fig. 3B**). Consistent with inhibited *RUNX1* and *THBS1*, expression of *CBFβ*, *CCL2*, *Cartilage Oligomeric Matrix Protein* (*COMP)*, *TGF-B1, ACTA2, COL4A1*, and *FN1* was attenuated in 5 out of 7 patients that received high-dose fresolimumab (**Supp Fig. 3C**). Collectively, these data imply that TGF-β signaling regulates the expression of *RUNX1* in SSc skin.

**Figure 3.**
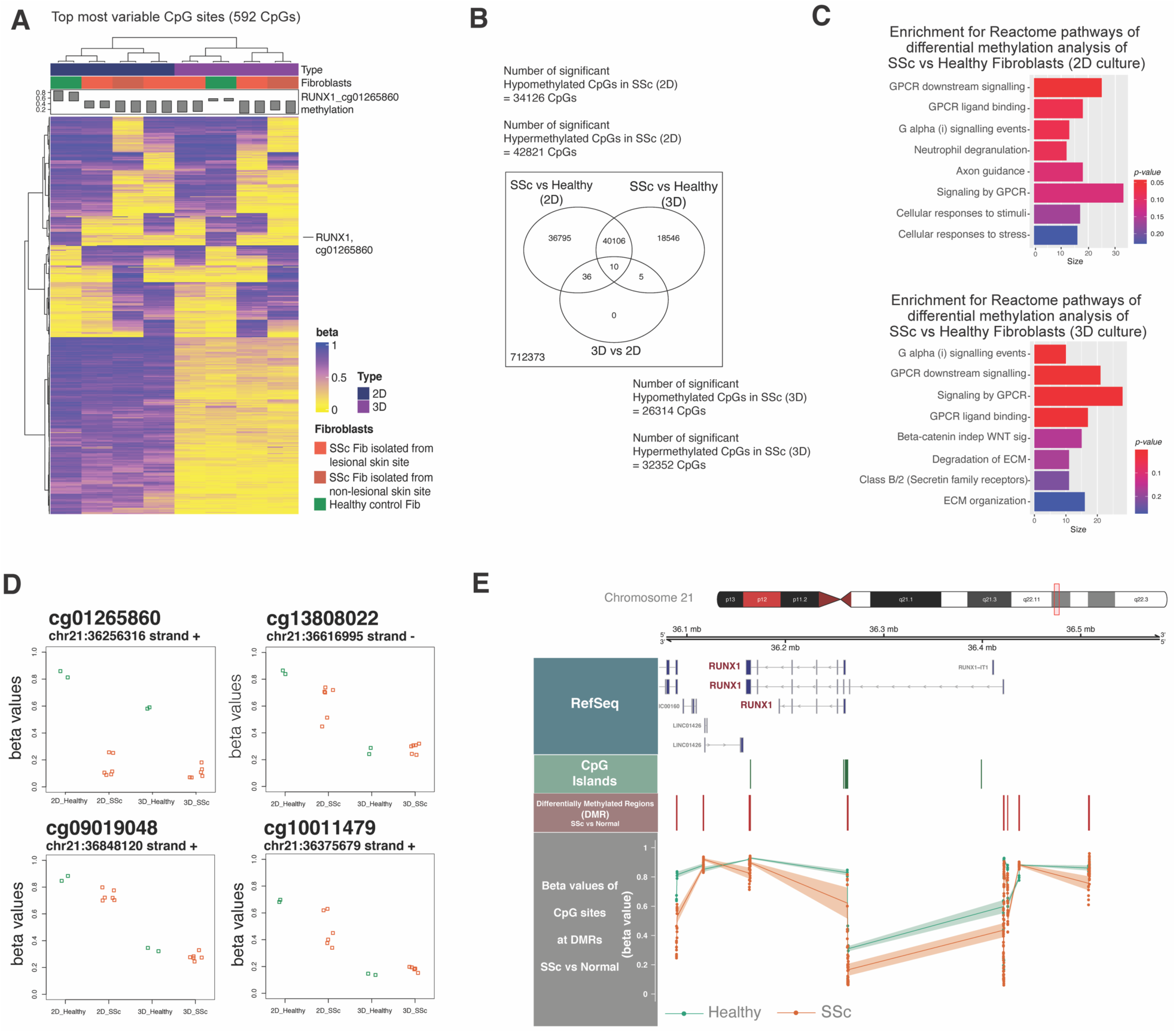
*RUNX1* is hypomethylated in SSc fibroblasts. **(A)** DNA methylation profile of 2D- and 3D-cultured fibroblasts isolated from dcSSc patients or healthy donors, created using Illumina’s Infinium Methylation EPIC array. Heatmap shows top 592 methylated CpG sites, with blue/yellow gradient of beta values. The bar-plot on top shows *RUNX1* beta value that is labeled within the heatmap, showing that *RUNX1* is hypomethylated in dcSSc samples. **(B)** Result of paired-wise differentially methylated CpGs and the number of significant CpGs in each group. **(C)** Pathway enrichment analysis of Reactome gene sets using top significant CpGs identified in (B) for each 2D and 3D culture. **(D)** The beta values of representative CpGs in *RUNX1* locus in 2D and 3D SSc and healthy conditions. **(E)** *RUNX1* locus on chromosome 21 and common CpG islands in green. The differentially methylated regions (DMRs) are identified between SSc and healthy samples are shown in red. The beta values corresponding to the CPGs at DMRs for SSc (in orange) and healthy (in green).

### RUNX1 inhibition reduces fibroblast proliferation and contraction

The compound Ro5-3335 has been reported to directly interact with RUNX1 and its heterodimeric partner CBFβ, repressing RUNX1/CBFβ-dependent transactivation in reporter assays^30^. Therefore, we hypothesized that Ro5-3335 could attenuate SSc fibroblast activation and contraction by inhibiting RUNX1 activity. To test this, we treated TGF-β1-activated SSc fibroblasts with Ro5-3335. As shown in **Fig. 2E**, *TGF-β1* levels significantly decreased upon RUNX1 inhibition. Furthermore, Ro5-3335 signaling reduced the proliferation rate of fibroblasts over the course of three days (**Fig. 2F**). Inhibition of RUNX1 activity by Ro5-3335 also significantly reduced the ability of normal human dermal fibroblasts to contract collagen gel matrices (**Fig. 2G, 2H**). Collectively, these results demonstrate that inhibition of RUNX1 activity reduced *TGF-β1* expression, as well as fibroblast cell proliferation and contractility in a collagen gel matrix, suggesting that RUNX1 is a driver of pathogenic SSc fibroblasts.

### Epigenetic dysregulation of RUNX1 in SSc fibroblasts

To evaluate the epigenetic state of *RUNX1* in SSc fibroblasts, we conducted genome-wide DNA methylation profiling using Illumina’s Infinium Methylation EPIC array to characterize fibroblasts in either 2D culture or in a 3D self-assembled (SA) skin-like tissue model culture^31,32^. The clinical characteristics of the dcSSc patients and healthy doners who provided samples for this experiment are provided in **Supp Tables S3** and **S4**. Two skin biopsies were collected from each donor, with one biopsy taken from forearm skin (lesional) and the other from flank skin (non-lesional). DNA was extracted from the isolated fibroblasts that were cultured on cell culture plates (2D culture) or in a 3D SA tissue on Millicell cell culture insert plates (3D culture). The methylation EPIC array was used to assess cytosine methylation at over 850,000 CpG sites across the genome. Once processed, data were filtered based on the most variated methylated sites; 592 CpG sites passed this filter and were hierarchically clustered by beta values (**Fig. 3A, Supp Table S5**). The beta value shows the degree of DNA methylation at a specific genomic locus, where 0 represents completely unmethylated CpG sites and 1 represents completely methylated CpG sites.

Samples clustered based on the most variable probes and were grouped primarily by the type of culture (i.e., 2D or 3D) and then by the type of sample (i.e., SSc or healthy) (**Fig. 3A**). Interestingly, for one case, there were differences noted in several CpG sites when comparing fibroblasts from that donor’s lesional versus non-lesional skin samples. This suggested that anatomic, site-specific differences in methylation patterns exist in isolated cells. Next, to understand if the type of culture (2D vs. 3D) has a significant effect on SSc or healthy fibroblasts, we performed probe-wise differential methylation analysis between SSc and normal samples in each culture type using the ‘limma’ package. In 2D culture, 34,126 CpGs were found to be significantly hypomethylated and 42,821 CpGs were hypermethylated in SSc samples relative to healthy controls. In 3D culture, 26,314 and 32,352 CpGs were hypomethylated and hypermethylated, respectively, in SSc samples relative to healthy controls. Among all significant CpGs (hypo- or hyper-methylated), 40,106 were common between the 2D and 3D culture, which suggests that the culture system may affect cytosine methylation rates in fibroblasts (**Fig. 3B**). We recognize that these differences in CpG methylation, however, may not translate to altered gene regulation and pathway activity. Therefore, we used the top significant probes to analyze the enrichment of Reactome pathways^33^ using the methylGSA package, which accounts for the number of CpGs per gene. As shown in **Figure 3C**, there were significant pathway differences between SSc fibroblasts and healthy control cells in both 2D and 3D culture, while no significant pathway differences were found between the different culturing conditions.

Most importantly, through the analysis of the most highly variable CpG sites, we found that one CpG annotated at the *RUNX1* genomic locus (cg01265860, located on the positive/sense strand of ch21:36256316) is hypomethylated in all SSc fibroblasts, compared to healthy control fibroblasts. Moreover, we also found a number of additional probes—including cg09019048 and cg10011479 on the sense strand and cg13808022 on the antisense strand—that were located on the *RUNX1* locus and were also hypomethylated (**Fig. 3D**). Hypomethylation in a gene on the sense strand is often associated with increased gene expression. While methylation on the antisense strand may not directly impact the expression of the gene, it can play a regulatory role, potentially through effects on RNA stability or processing.

Next, we analyzed the differentially methylated regions (DMRs) between SSc and healthy fibroblasts to identify proximal CpGs that are concordantly differentially methylated (**Supp Table S6**). **Figure 3D** shows the *RUNX1* locus on chromosome 21 using the hg19 reference genome. The *RUNX1* gene is indicated and common CpG islands are shown (**Fig. 3E**). Several DMRs were evident between SSc and healthy control fibroblasts. Analysis of the average beta values of CPGs at DMRs surrounding *RUNX1* demonstrated that the SSc samples were hypomethylated at almost all the sites compared to healthy controls (**Fig. 3E**). Collectively, our data indicate that the significantly elevated high *RUNX1* expression observed in patients with SSc is most likely due to dysregulation at the epigenetic level.

### RUNX1 is expressed in a subpopulation of SSc-associated fibroblasts

To identify the fibroblast subpopulation(s) expressing *RUNX1*, we analyzed the publicly available single-cell RNA sequencing (scRNA-seq) data from 12 dcSSc patients and 11 matched healthy controls (GSE138669)^34^. The skin biopsies in this study were collected from dorsal mid-forearm and a single-cell suspension was created through enzymatic digestion^35^. **Figure 4A** shows the major cell types in a uniform manifold approximation and projection (UMAP) plot after data processing. The total *RUNX1* aggregate expression was then calculated and displayed in each sample. While heterogeneity exists between the patients, *RUNX1* was more highly expressed in dcSSc samples than in healthy control skin samples, in accordance with bulk gene expression analyses (**Fig. 4B**). Notably, the SSc patient with the highest *RUNX1* expression levels was treatment naïve with early-stage disease (duration of ∼10 months; donor ID: SC189).

**Figure 4.**
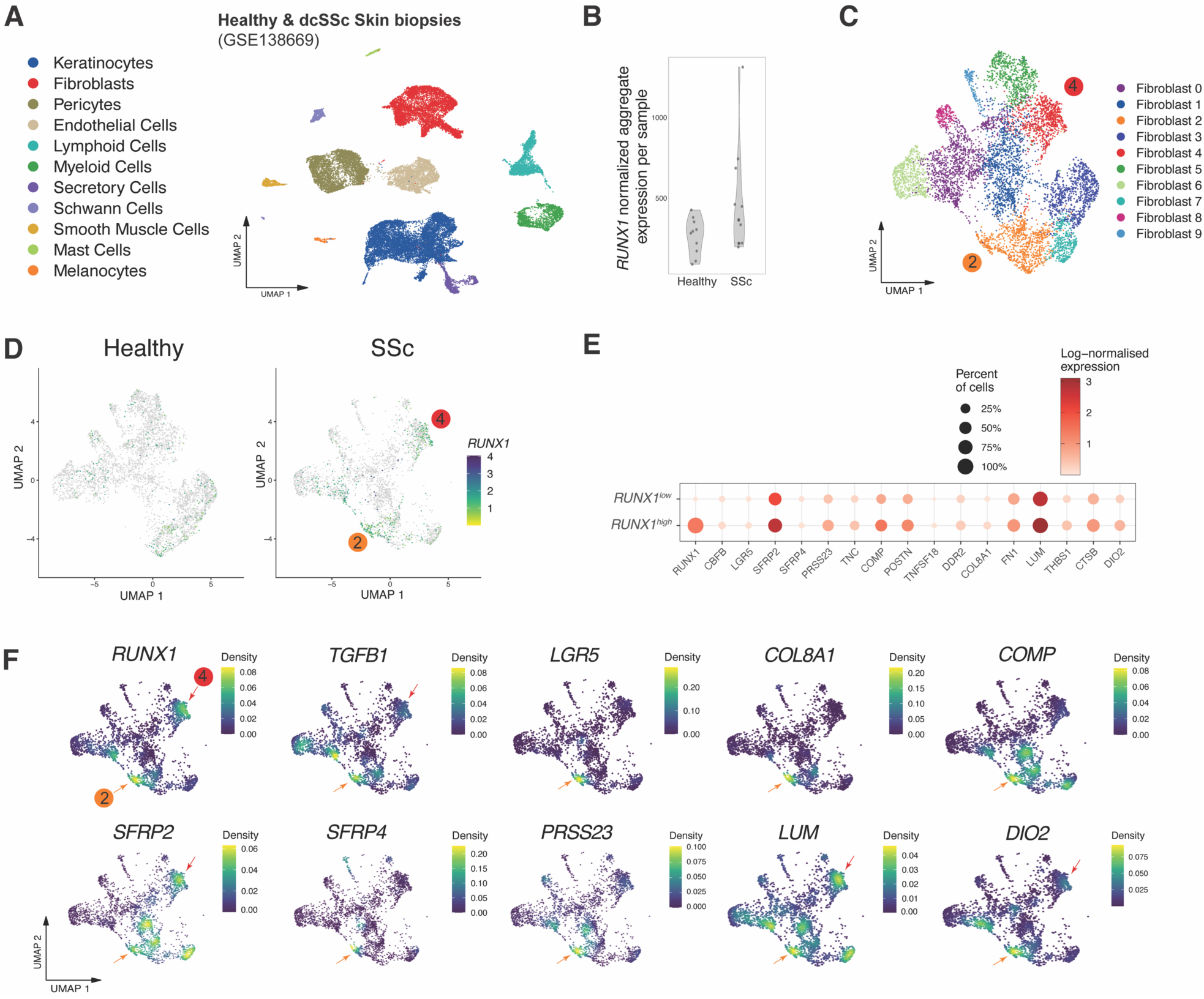
Enrichment of *RUNX1* in SSc-specific fibroblast subpopulation. **(A)** UMAP projection of cell types from Tabib et al., 2021’s scRNA-seq of forearm skin biopsies **(B)** *RUNX1*-normalized aggregate expression of 10 samples from healthy donors and 12 from dcSSc patients. **(C)** UMAP projection of 10 fibroblast subpopulations (clusters 0–9). Two fibroblast populations of 2 and 4 are marked, which are enriched SSc samples. **(D)** Feature plots of *RUNX1* expression in healthy and SSc fibroblasts. Two *RUNX1*-expressing fibroblast clusters are marked with their respective numbers. **(E)** The log-normalized expression rate of main differentially expressed genes between *RUNX1^high^*- with *RUNX1^low^*-expressing SSc fibroblasts. **(F)** Density plots of *RUNX1* and major SSc-relevant genes within SSc fibroblast subpopulations. Arrows indicate the cluster 2 and 4 of SSc-specific subpopulations of fibroblasts.

To examine *RUNX1* expression in the fibroblast subpopulations, the total fibroblast populations were clustered into 10 groups (clusters 0–9). As demonstrated in **Figs. 4C and 4D**, *RUNX1* was enriched in SSc-enriched subpopulations 2 and 4. Comparison of *RUNX1^high^*- with *RUNX1^low^*-expressing single cells showed that the fibroblasts with high *RUNX1* expression levels also highly expressed fibrotic genes, such as *FN1*, *LUM*, *POSTN*, *COMP*, *COL8A1*, and *TNC* (**Fig. 4E**).

Additionally, we observed that the *RUNX1* is expressed in *LGR5^+^* and *TGF-β1^+^* cells (**Fig. 4F**). *LGR5^+^* fibroblasts are associated with ECM degradation and skin remodeling/assembly genes^36^. Moreover, *RUNX1*-expressing fibroblasts are *SFRP2^+^/SFRP4^+^*, which have been shown to be associated with the myofibroblast signature^35^.

## Discussion

Efforts to fully characterize the driving factors of SSc dermal fibroblast heterogeneity and translational states are limited by tissue scarcity, anatomic site-specific differences, and a paucity of reliable fibroblast subset markers^37^. Defining SSc-specific fibroblast regulatory programs that drive fibrosis or, conversely, promote regeneration, is essential for achieving favorable healing outcomes. Here, we analyzed SSc patient heterogeneity in terms of *RUNX1* expression in multiple cohorts and established a more comprehensive understanding of *RUNX1* in SSc. We demonstrated that *RUNX1* is enriched in both lesional and non-lesional skin, with highest expression in dcSSc patients. As non-lesional skin samples may yield information about the underlying disease process in most skin inflammatory diseases^28^, the observation of *RUNX1* abnormalities in both lesional and non-lesional skin indicates that *RUNX1* may be a driver of SSc pathogenesis in a pre-lesional state. Additionally, we have demonstrated that higher expression of *RUNX1* is correlated with increased disease severity in patients with SSc, as evidenced by a greater risk for elevated mRSS and the presence of ILD.

Konkimalla *et al.* have recently shown that *RUNX1* is a key driver of alveolar fibroblast transitional states in multiple lung-injury mice models, and that loss of *Runx1* leads to a decrease in the pulmonary levels of ECM components and their assembly^38^. The mechanisms by which RUNX1 controls the expression of ECM components in TGF-β-activated fibroblasts is not yet clear. TGF-β signaling is known to enhance fibrogenesis and stimulate the production of ECM components, including collagen, fibrillin, laminin, fibronectin, proteoglycans, and elastin^39^. Previous studies have revealed that RUNX1 is essential for mesenchymal stem cell proliferation and myofibroblast differentiation, and associates with the SMAD-dependent TGF-β signaling pathway to directly bind promoters and enhancers of multiple cell-state-specific genes related to cell proliferation and migration^8,40,41^. Similarly, we demonstrated that the inhibition of RUNX1 via Ro5-3335 reduced dermal fibroblast proliferation and contraction, further supporting the hypothesis that RUNX1 is a driver of fibroblast activation.

*RUNX1* upregulation is a hallmark of several other fibrotic diseases. For example, *RUNX1* is expressed in human retinal microvascular endothelial cells in proliferative vitreoretinopathy (PVR)^42^. In agreement with our findings, inhibition of RUNX1 activity resulted in a significant reduction of retinal lesion size and reduced migration, proliferation, and tube formation of human retinal microvascular endothelial cells *in vitro*^42–44^.

We have also demonstrated that *RUNX1* is hypomethylated in the SSc-derived fibroblast genome, indicating that the increased *RUNX1* expression observed in SSc patients is likely due to dysregulation at the epigenetic level. Using a single-cell multiomic analysis, in a separate study, we showed that *RUNX1* is consistently enriched in a subpopulation of fibroblasts in an SSc-derived self-assembled skin-equivalent (saSE) tissue model at levels similar to those observed in SSc skin biopsies^45^. In an analysis of single-cell sequencing assay for transposase-accessible chromatin (scATAC-seq), a fibroblast subpopulation was identified by elevated TF motifs for *RUNX1* and *RUNX2*, together with *SMAD3*, *SMAD5*, and *NFKB1*. This population exhibited increased expression of multiple collagens and pro-fibrotic genes, including *POSTN* and *LUM*. In light of DNA methylation and enriched chromatin accessibility data, which indicate higher TF motif accessibility, these observations imply that SSc fibroblasts are more susceptible to activation and differentiation under the influence of RUNX1.

Recent work using scRNA-seq from SSc skin biopsies demonstrated the existence of distinct dermal fibroblast subsets that are specific to SSc^34,46,47^. Here, we re-analyzed publicly available data from Tabib *et al.* for altered gene expression in SSc-enriched fibroblast populations. An increased proliferation of *SFRP2^hi^PRSS23^+^* fibroblasts was identified in SSc skin, which is thought to represent a progenitor population for myofibroblasts. Interestingly, this population of fibroblasts has *RUNX1^high^* activity. Notably, dysregulation of associated genes in the *Wnt* signaling pathway, *SFRP2* and *SFRP4* (*Secreted Frizzled-Related Protein 2 and 4*), has been implicated in the pathogenesis of fibrosis and metabolic disease^48,49^.

Another scRNA-seq and scATAC-seq study that included a larger population of SSc patients identified a distinct SSc-specific subpopulation of fibroblasts termed as *LGR5^+^-*scleroderma-associated fibroblasts (*LGR5^+^-*ScAFs)^36^. In dcSSc patients, *LGR5^+^-*ScAFs exhibited increased expression of genes associated with ECM components such as collagens, *POSTN*, and *COMP*. Additionally, genes such as *PRSS23*, *SERPINE2*, *TGFB*, *SFRP4*, *DKK3*, *THBS1* and *CTGF* were upregulated in ScAFs^36^. Our data demonstrated that *RUNX1^high^* expressing fibroblasts are *LGR5^+^* cells, suggesting that *RUNX1* is an SSc fibroblast TF that contributes to the altered phenotype of these cells.

A recent study of scRNA-seq and spatial transcriptomics analyses of SSc skin by Ma *et al.* identified two major clusters of *SFRP^+^* fibroblasts and *COL8A1^+^*fibroblasts^50^. *COL8A1^+^* fibroblasts were located in the deeper dermis of SSc skin and expressed high levels of *ACTA2*, *SFRP4*, *POSTN*, and *PRSS23*, and also had the highest ECM score. Notably, we show here that *RUNX1* is activated in *COL8A1^+^* cells. While variation exist among the fibroblast subpopulations described in patient cohorts of these different studies, which may arise from different RNA sequencing depth and the type of library preparation method, the expression of *RUNX1* is consistently linked to either the myofibroblast or SSc-specific population, or the direct progenitors of these cells in each of these cohorts.

Leveraging gene expression, DNA methylation, and single-cell resolution data of SSc fibroblasts, we discovered unique associations between disease-specific subpopulations of fibroblasts and *RUNX1*, which may serve as a potential novel therapeutic target for SSc. As *RUNX1* activity is tissue-specific, our study is limited by identification of the direct target genes in SSc fibroblasts. In this regard, ChIP-seq (chromatin immunoprecipitation sequencing) studies may be used to identify DNA-binding sites for RUNX1 and to infer potential target genes in SSc-relevant cell types. The recent RUNX1 ChIP-seq study in glioblastoma identified ECM genes such as *COL4A1*, *LUM*, and *FN1* as targets^18^. It is possible that these genes are similarly regulated by RUNX1 in dermal fibroblasts. Future studies focused on profiling of *RUNX1* deletion or inhibition in SSc fibroblasts will not only provide insights into the direct cellular targets and involved pathways but may also inform whether the inhibition of RUNX1 is sufficient to block progression and reverse established SSc dermal fibrosis.

## Materials and Methods

### DNA microarray data processing

#### Agilent DNA microarray processing

Raw gene expression data from skin biopsies of patients with SSc and healthy controls were obtained from the gene expression omnibus (GEO) including: Hinchcliff *et al.*, 2013^25^ (GSE59787, Fig. 1), Milano *et al.*, 2008^3^ (GSE9285, Supp Fig. 2A), Pendergrass *et al.*, 2012 ^23^ (GSE32413, Supp Fig. 2B), Franks *et al.*, 2019 (GSE125362, Supp Fig. 2C), Gordon *et al.*, 2018 (GSE97248, Supp Fig. 2D), and one database with TGF-β1-induced fibroblasts^29^ (GSE12493, Fig. 2). Information on all of the datasets is provided in **Supp Table S1**. The clinical data were either publicly available or collected upon request from the authors.

Samples were assayed using Agilent Technologies 2-channel DNA microarrays platform: Agilent-014850 Whole Human Genome Microarray 4x44K G4112F platform, or Agilent-012391 Whole Human Genome Oligo Microarray G4112A platform, and Agilent-028004 SurePrint G3 Human Gene Expression 8x60K Microarray platform. Raw data in the format of the .gpr file generated by GenePix scanner were processed through Bioconductor ‘*limma’* package by reading the Cy3/green channel of the arrays (mean foreground and median background signal intensities). Low-quality spots were weighted, and outlier spots were identified using median absolute deviation (∼0.6% spots were found to be outliers based on the quality). The method of normexp^51^ was used for background correction followed by quantile normalization to enable cross-array comparisons. Probes with >80% of 1.3-fold or lower intensity over local background were considered as very-low-expressed probes within each data set and were removed. The Agilent’s annotations were downloaded from “Agilent Technology eArray” webpage for each of the microarray platforms. Using these annotations, probe IDs were then converted to gene symbols and ‘*collapseRows’*^52^ was used to collapse the normalized log2 intensities based on gene symbols. Finally, the normalized log2 expression matrices were used to measure the expression rate of *RUNX1* in all arrays at different time points.

The Hinchcliff *et al.*, 2013 (GSE59787) dataset includes skin biopsies from longitudinal data with samples from baseline and at 6, 12, 24, and 36 months from both lesional and non-lesional skin. In all databases, the existence of batches was evaluated before the analysis and corrected for. Patients who had been diagnosed with the disease within 2 years of the date of skin biopsy were assigned as “early stage” and the rest were “late stage.” For Pendergrass *et al.*, 2012 (GSE32413) only samples from one batch at baseline were used. Gordon *et al.*, 2018 (GSE97248) had samples from baseline and 12 months; therefore, both time points were analyzed. In the TGF-β1-induced fibroblast^29^ (GSE12493) dataset, two separate platforms were used (Agilent-014850 and Agilent-012391). As a result, data were first analyzed separately under each platform and then carefully merged by ‘ComBat’ from ‘sva’ package (v. 3.46.0) with a design matrix to control for biological differences between clinical conditions while performing batch effect correction.

#### Illumina DNA microarray processing

Raw microarray data were obtained from GEO using accession ID GSE58095 (Assassi *et al.*, 2015^27^). The data were generated on the Illumina HumanHT-12 V4.0 expression beadchip platform and processed using the ‘lumi’ package (v. 2.50.0) in R (v. 4.2.2). Probes with high detection (*P*-values > 0.01) were removed to filter out probes with low signal-to-noise. The ‘arrayQualityMetrics’ package (v. 3.54.0) was utilized to perform quality control assessment of the processed data. The ‘illuminaHumanv4’ was employed to annotate the probes. This package provides annotation information for probes, including gene symbols and genomic coordinates. Normalized and transformed expression values were then obtained using quantile method by ‘*normaliseIllumina’* implemented in the ‘*beadarray’* package. Similar to the Agilent pipeline, we used ‘*collapseRows’* to collapse the intensities based on gene symbol.

#### Affymetrix DNA microarray processing

Raw microarray data were obtained from GEO using accession ID GSE55036 (Rice *et al*., 2015). The data were Affymetrix DNA microarrays that were processed using the ‘Affy’ package (v. 1.76.0). The ‘gcrma’ algorithm was utilized to convert the raw data into an ExpressionSet object. This algorithm employs robust multi-array averaging (RMA) with additional sequence-based correction to estimate expression values. Quality assessment was performed using the Relative Log Expression (RLE) and Normalized Unscaled Standard Error (NUSE) metrics. These metrics assess the distribution of expression values across samples and provide insights into the overall data quality and consistency. The ‘*arrayQualityMetrics*’ package was used to assess various aspects of microarray data quality. Based on these QC assessments, three low-quality arrays from patient 19 (at 3, 7, and 24 weeks) were removed from subsequent analysis. The appropriate annotation package of ‘*hgu133a2.db’* (v. 3.13.0) and ‘*collapseRows’* was utilized to annotate and convert the probes to gene symbols.

### Differential expression analysis (DEG)

Differential expression analysis was performed on the processed microarray data using the ‘*limma*’ package. Contrast of interest between 12 hours after exposure and baseline was specified using a contrast matrix. Significant DEGs were annotated and further analyzed to interpret biological significance using Reactome gene sets.

### Gene Set Variation Analysis (GSVA)

The resultant normalized log2 expression matrices of GSE59787 and GSE55036 were subsequently employed in Gene Set Variation Analysis (GSVA). GSVA is a non-parametric method for estimating variations in gene set enrichment within a given set of samples derived from an expression dataset^53^. The ‘*gsva*’ (v. 1.51) algorithm was used to calculate the samples’ enrichment scores for cell-type signatures across all samples. The gene sets are listed in **Supp Table S2**.

### Fibroblast isolation and culture

SSc and healthy dermal fibroblast lines were isolated and expanded from skin punch biopsies as described previously^31,54,55^. Isolated fibroblasts were stored in liquid nitrogen in stocks of 0.5 million cells in freezer media [70% fibroblast media, 20% FBS (fetal bovine serum, HighClone), and 10% dimethyl sulfoxide (DMSO)]. Fibroblasts were maintained in complete growth media [90% DMEM (Dulbecco’s Modified Eagle Media), 10% FBS, 1% HEPES (Millipore Sigma, Germany)], and 1% penicillin-streptomycin (Corning). Fibroblasts at passages less than 8 were used to seed for 2D and 3D tissue cultures. A summary of demographic and clinical characteristic of the patients and healthy controls who donated skin biopsies for DNA methylation profiling is provided in **Supp Table S3**. For the RUNX1 inhibition assay, TGF-β1-induced (10 ng/mL) (R&D systems, #240-B) SSc fibroblasts were treated with 20 μM Ro5-3335 (TOCRIS, #4694).

### Quantitative real-time PCR

Total RNA was isolated using the RNeasy Fibrous Tissue Mini Kit (Qiagen, 74704) per manufacturer’s instructions. Complementary DNA (cDNA) was synthesized from 60 ng total RNA using the qScript™ Ultra cDNA SuperMix (QuantaBio, #95161). Quantitative real-time PCR was performed using TaqMan Universal PCR Master Mix (Life Technologies, #4324020) for human *ACTB* (Applied Biosystems, #Hs01060665_g1), and *TGF-β1* (Applied Biosystems, #Hs00998133_m1). The StepOnePlus Real-Time PCR System (Applied Biosystems) was used with threshold cycle number was determined using Opticon software. The mRNA levels were °threshold for the control gene; Et denotes the average threshold for gene of interest.

### Collagen contraction assay

Measurements of 3D fibroblast collagen contraction were performed as previously described^56^. Briefly, culture plates are pre-coated with BSA and one part 4x10^4^ fibroblasts/mL suspension in MCDB medium (Sigma, #M6395) mixed with collagen solution, which is a mix of one part collagen (Advanced BioMatrix, 6 mg/mL), one part HEPES (pH 8), and two parts MCDB 2X. Then, 1 mL of collagen suspension is added to the pre-coated wells and allowed to polymerize. The final collagen concentration is 1.2 mg/mL with 80,000 cells/mL. For fixed contraction assays, cells were detached from the wells 48 hours post-polymerization and contraction was quantified by a decrease in gel diameter within 5 hours. For floating assay, the cells were detached from the plate immediately after polymerization and the gel diameter was measured after 48 hours in a 37°C incubator. Cells were pre-incubated with Ro5-3335 and SIS3 (TOCRIS, #5291, as positive control) for one hour prior to the assay. In principle, contraction in the floating model occurs in the absence of external mechanical load, whereas, in the fixed model, the attachment of the cell-collagen mixture to the tissue culture dish induces more stress fibers.

### DNA extraction, bisulfite conversion, and DNA methylation array

A total of 16 samples [eight samples from 2D monolayer cultures of fibroblasts isolated from patients with SSc or healthy controls, and eight samples from 3D cultures (2 replicates from each biological sample)] were selected for DNA extraction. Genomic DNA was extracted using the Qiagen QIAcube by QIAamp DNA Mini Kit (Hilden, Germany). To ensure that all samples met DNA quality control criterion, we used Agilent 4200 TapeStation (Genomic DNA ScreenTape, Agilent technologies, Santa Clara, CA). Between 500 ng and 1 µg of DNA from these 2D and 3D samples were bisulfite converted and processed according to the Illumina Infinium MethylationEPIC array protocols at the University of Southern California’s Molecular Genomics Core Laboratory (Los Angeles, CA).

### DNA methylation data processing

The raw intensity data (IDAT files) from the MethylationEPIC array were processed using the ‘minfi’ package (v. 1.44.0), which provides functions for reading, preprocessing, and analyzing DNA methylation data. We filtered the probes with *detection P-values* < 10^-5^, which compares the total, methylated (M), and unmethylated (UM) signals for each probe to the background signal level. Thereby, probes with high *detection P-values* or high background noise were excluded from further analysis. The data were then normalized using quantile normalization.

Probes with CpGs at common single nucleotide polymorphisms (SNPs) or that tracked to sex chromosomes were filtered out, leaving a total of 807,871 probes. The ‘*Illumina Human Methylation EPICanno.ilm10b4.hg19’* (v. 0.6.0) package was used to annotate the CpG probes. This package provides annotation information for probes, including corresponding gene reference, genomic coordinates, type of strand, and probe sequence. Finally, methylation levels were calculated through both beta values (β=M/(M+UM)) and M-values (M-value=log2(M/UM)), which were used for downstream analyses. Topmost variable probes were used for unsupervised hierarchical clustering to examine how the samples cluster with each other using beta values. Additionally, probe-wise differential methylation analyses were performed using the ‘*limma’* package on M-values with a design matrix accounting for SSc vs. healthy conditions in 2D and 3D culture systems. The most significant differentially methylated CpGs were subsequently used for gene set enrichment analysis by the ‘*methyglm’* function from the ‘*methyGSA’* package; this carries out gene set analysis adjusted for the number of CpGs per gene.

### Differential methylation analysis of regions (DMRs)

The ‘*DMRcate*’ package was used to identify differentially methylated regions (DMRs) between SSc and healthy samples using M-values, which are proximal CpGs of genes that are concordantly differentially methylated between the conditions. The DMRs corresponding to the *RUNX1* gene are provided in **Supp Table S6**.

### Processing of single-cell RNA sequencing

Single-cell RNA-seq data from Tabib *et al.* was downloaded from GEO: GSE138669. Raw sequencing data were initially assessed through *Cell Ranger*, v. 6.0.1. The High-Performance Computing (HPC) resources at Dartmouth College were used to integrate the data. For data processing, the ‘*Seurat*’ package v. 4.4.0 was used. Quality control steps were implemented to filter out low-quality cells based on low counts, high mitochondrial gene expression, and doublet detection criteria. Cells were filtered with unique feature counts between 200–2,500 and less than 5% mitochondrial reads. It should be noted that data was of low-read depth (majority of cells <50 features per cell), so a large number of cells were lost through filtering steps. Our quality control filtering resulted in a total of 33,171 single-cell transcriptomes.

The count matrix was normalized using the ‘*NormalizeData*’ method implemented in ‘Seurat’. Additionally, gene expression values were scaled across cells, using the ‘*ScaleData*’ function. Normalized and scaled expression values were then used for downstream analyses. Cell type was assigned to each cluster using gene expression distribution of cell-type-specific markers shown in each original publication and using ‘*FindNeighbors*’ and ‘*FindClusters*’ functions.

To identify fibroblast clusters based on similar gene expression profiles, the ‘*FindClusters*’ were applied only to fibroblast clusters using a graph-based clustering approach. A total of 10 clusters were found, consistent with the original publication.

To calculate the total *RUNX1* expression on each sample, ‘*AggregateExpression*’ was performed on the log normalized data. The library of ‘*scCustomize*’ was used to make UMAP plots, feature plots, and density plots. The ‘*FindMarkers*’ function was used to identify the top genes between *RUNX1^high^* and *RUNX1^low^*fibroblast populations.

## Statistical analyses

Statistical significance was determined using a Student’s *t-test*, paired *t-test*, or Pearson correlation test, as appropriate, and a *P*-value of <0.05 was considered significant. All data were analyzed using R (v. 4.2.2). For gene sets, effect size measurements of Hedge’s g were used to quantify the standardized mean difference between two groups for each gene set. Hedge’s g effect size is a measure of association and not a measure of statistical significance.

## Data availability

The raw IDAT files for SSc and healthy DNA methylation will be deposited into the Gene Expression Omnibus (GEO) and will be made publicly available. Any additional data and R analysis files utilized in this study may be obtained from the lead contact upon request.

## Authors’ contributions

Conceptualization: RP, PAP, MLW

Data Analysis: RP, ZG, HCJ

Experimental assays: RP, DMT, DP, SS

Investigation: RP, TRA, ZG, DMT

Funding Acquisition: JG, MLW

Supervision: MLW

Writing-original draft: RP

Writing-review and editing: RP, ZG, HCJ, MJM, TRA, DP, PAP, MLW

## Acknowledgements

For advice on data and statistical analysis, we would like to thank the Dartmouth Data Analytics Core. The authors also acknowledge the support of the Laboratory for Clinical Genomics and Advanced Technology in the Department of Pathology and Laboratory Medicine of the Dartmouth. Finally, we thank our funding sources, including the National Institute of Allergy and Infectious Diseases (NIAID) R21AI169420 (PAP and MLW), and the National Institute of Arthritis and Musculoskeletal and Skin Diseases (NIAMS) R21AI178651 (PAP and MLW).

## Supplementary Figures

**Supp Figure 1.**
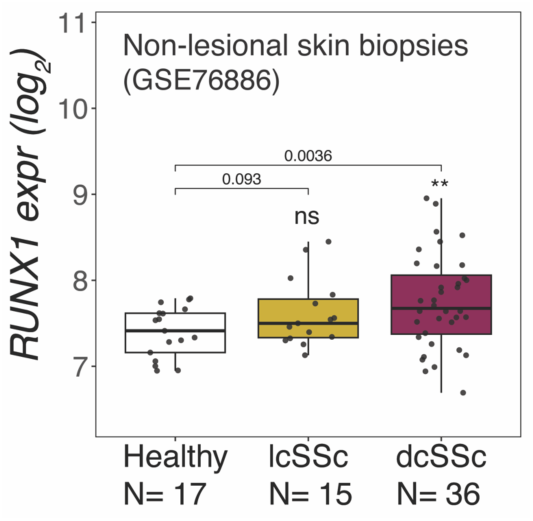
*RUNX1* expression rate in flank (non-lesional) skin biopsies of dcSSc patients (*N*=36), lcSSc patients (*N*=15), and healthy donors (*N*=17).

**Supp Figure 2.**
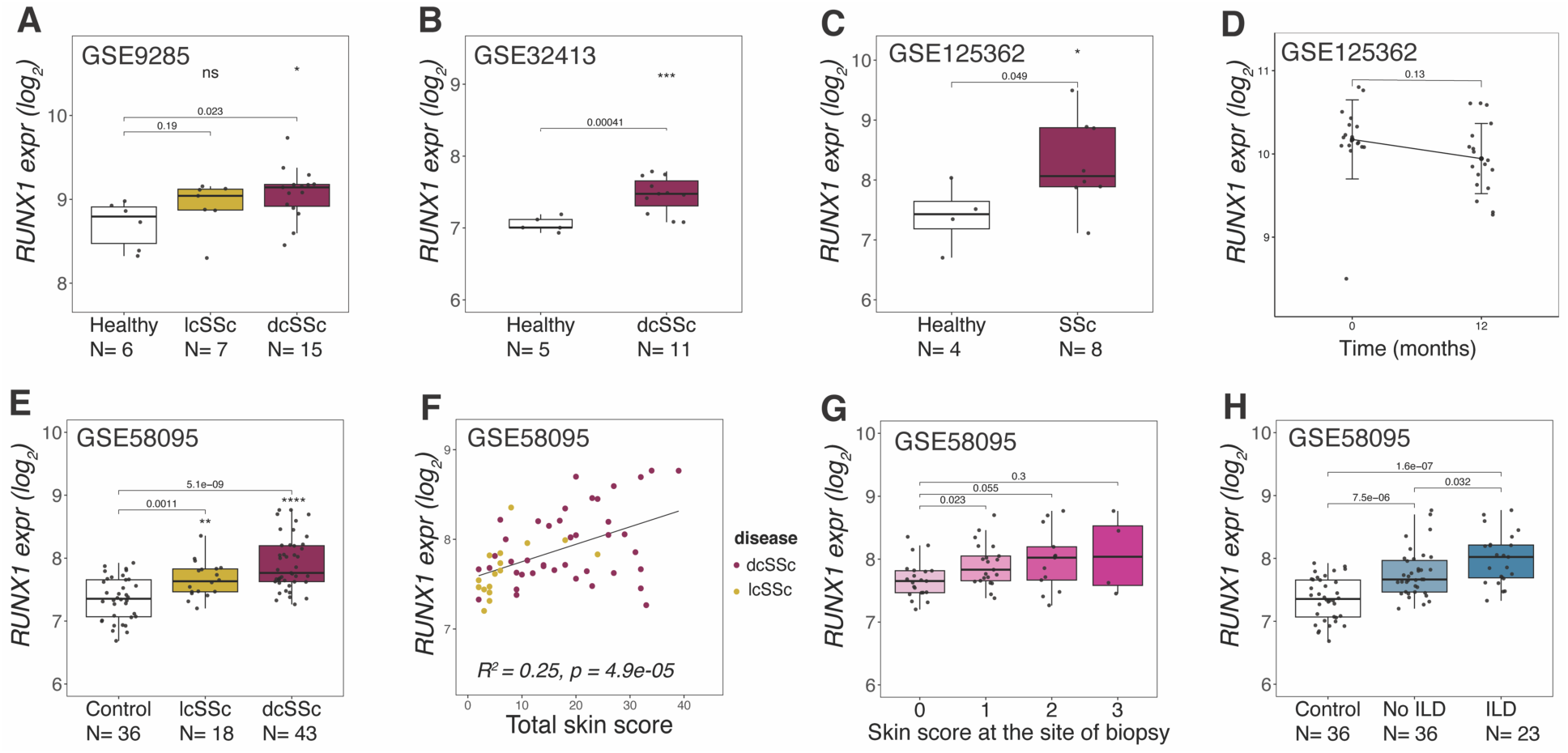
(A) *RUNX1* expression rate in skin biopsies of dcSSc patients (*N*=15), lcSSc patients (*N*=7), and healthy donors (*N*=6) from the Milano *et al.*, 2008 study (GSE9285). (B) *RUNX1* expression rate in skin biopsies of dcSSc patients (*N*=11) and healthy donors (*N*=5) from the Pendergrass *et al*., 2012 study (GSE32413). (C) *RUNX1* expression rate in skin biopsies of dcSSc patients (*N*=8) and healthy donors (*N*=4) from the Franks *et al.*, 2019 study (GSE125362). **(D)** *RUNX1* expression rate in skin biopsies of dcSSc patients (*N*=18) at baseline and a 12-month follow-up from the Gordon *et al.*, 2018 study (GSE97248). **(E)** *RUNX1* expression rate in skin biopsies of dcSSc patients (*N*=43), lcSSc patients (*N*=18), and healthy donors (*N*=36) from the Assassi *et al.*, 2015 study (GSE58095). **(F)** Correlation between *RUNX1* expression and mRSS skin score at baseline for both lcSSc (yellow) and dcSSc (red) patients. **(G)** *RUNX1* expression rate to the skin score at the site of biopsy. **(H)** *RUNX1* expression rate in healthy donors (*N*=36), patients with ILD (*N*=23), and patients without ILD (*N*=36) at the time of biopsy.

**Supp Figure 3.**
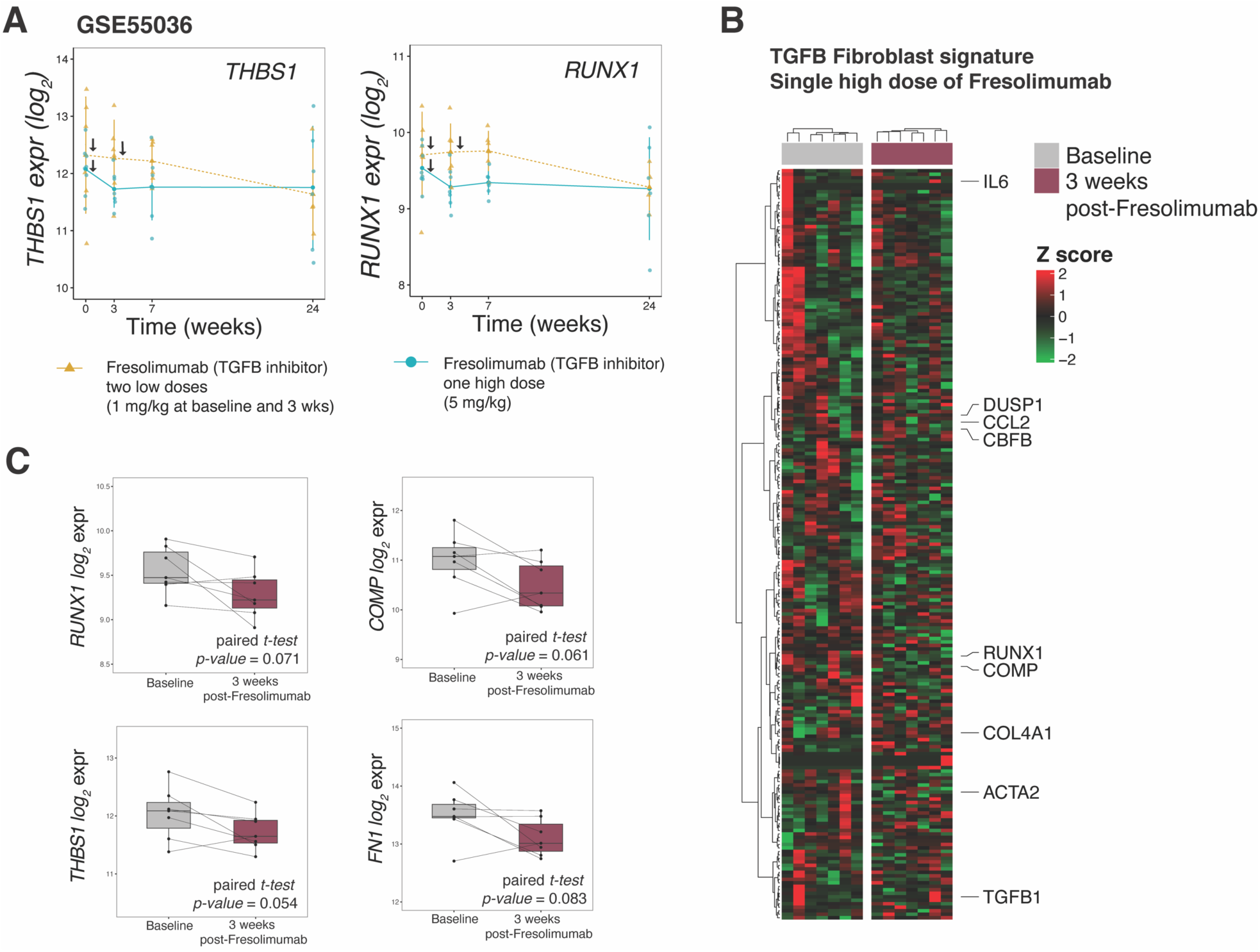
The TGF-β inhibitor fresolimumab reduces *RUNX1* expression in SSc skin. **(A)** *THBS1* and *RUNX1* expression levels in dcSSc skin biopsies of patients who were given two low doses (1 mg/kg) of fresolimumab at weeks 1 and 3 in yellow (*N*=7); or a single high dose (5 mg/kg) of fresolimumab at week 1 in blue (*N*=7). The mid-forearm skin biopsies were collected at baseline and again at weeks 3, 7, and 24. (B) The heatmap of genes in the TGF-β fibroblast cell signature for patients who received a high dose of fresolimumab at baseline and again 3 weeks after treatment (*N*=7). (C) The expression of several genes including *RUNX1* and TGF-β pathway biomarkers such as *COMP*, *THBS1*, and *FN1*.

## Notes

### Competing Interest Statement

MLW has received consulting and research funding from Bristol-Myers Squibb, Boehringer Ingelheim, Corbus Pharmaceuticals, Celdara Medical, LLC, and UCB Biopharma for systemic sclerosis research. All other authors declare no competing interests.

